# Unravelling the molecular activation of the reparative cardiac fibroblasts after myocardial infarction

**DOI:** 10.1101/2024.11.21.624638

**Authors:** Silvia C Hernández, Marina Ainciburu, Laura Sudupe, Nuria Planell, Amaia Vilas-Zornoza, María López-Moreno, Sarai Sarvide, Luis Diaz-Martinez, Jorge Cobos-Figueroa, Patxi San Martin-Uriz, Emma Muinos-López, Gloria Abizanda, Purificación Ripalda-Cemboráin, Vincenzo Lagani, Juan P. Romero, Jesper Tegner, José Mª Pérez-Pomares, Ming Wu, Stefan Janssens, Felipe Prósper, Adrián Ruiz-Villalba, David Gómez-Cabrero

## Abstract

Activated cardiac fibroblasts (*Postn*^+^ CFs) are responsible for the healing of the heart tissue after a myocardial infarction (MI). However, so far little is known about the moment that CFs are activated, and the genes involved in this process. This is especially relevant in the context of CF heterogeneity and their role in the response to the damage. In this context, we have described a subpopulation of activated CFs responsible for the healing scar and for preventing the rupture of the ventricle after the damage: the Reparative Cardiac Fibroblasts (RCFs). Our new data indicate that RCFs directly derived from activated CFs, and this transcriptional shift happens in a close window after damage. Interestingly, our results exhibited two different molecular dynamics that would give rise to this activation and, consequently, the appearance of definitive RCFs. Using bulk RNA-Seq, RNAScope and Spatial Transcriptomics, we anatomically localized some of the genes related to both dynamics in the infarcted heart and highlight the potential role of *Aspn* as a new marker of this transcriptional transition in mice, pigs and patients.

## Main text

Fibrosis can be considered as an evolutionarily conserved adaptative process in response to damage in both homeostatic and pathological contexts [1]. This process is characterized by remodeling of the extracellular matrix architecture and is led by fibroblasts. Although fibroblasts have been historically considered as a rather homogeneous cell population, recent studies have demonstrated their heterogeneity [2, 3]. Understanding this heterogeneity is key to unraveling the cellular and molecular mechanisms that drive the fibrotic process and thus modulate it.

Recently, we described, among 13 subpopulations, a subpopulation of activated cardiac fibroblasts (CFs) (*Reparative Cardiac Fibroblasts*, RCFs) as the primary driver of the healing process in the context of myocardial infarction (MI) [4]. We have hypothesized that *Cthrc1*^*+*^ RCFs represent the final activation stage of activated (*Postn*^+^) CFs. Interestingly, recent results in early phases of the fibrotic process in the lung revealed a subpopulation with a similar transcriptomic profile and function [3]. Based on this, defining the timing of RCF activation might be of importance for the development of personalized strategies to manage the initial phases of the repair process. In order to translate this knowledge into candidate therapeutic strategies, we studied the regulatory transcriptional dynamics governing RCF behavior.

To this end, we performed bulk RNA-seq analysis of isolated *Col1α1*-GFP^+^/CD31^-^/CD45^-^ CFs between 1- and 6-days post-infarction (dpi). Additional data from healthy, 7, 14 and 30 dpi CF samples were included in our study [4]. The expression of highly relevant RCF marker genes (*Cthrc1, Ddah1, Postn, Fn1, Lox* and *Ptn*) peaked between 3 and 5 dpi (Figure [A]). Accordingly, CTHRC1^+^ RCFs were identified in the infarcted tissues at these two time points (Figure [B]).

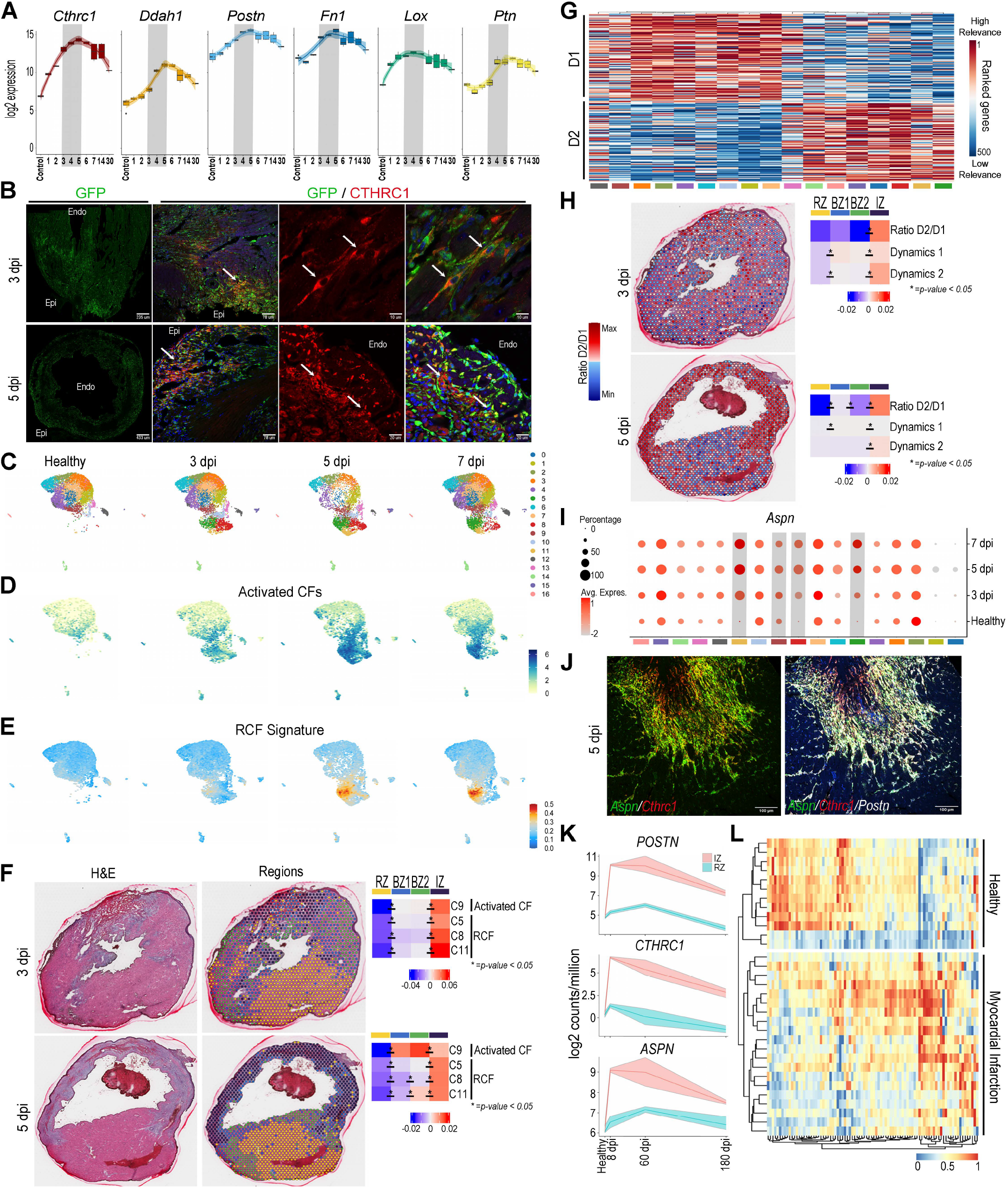

Next, to elucidate how *Cthrc1*^*+*^ RCFs are determined among *Postn*^+^ CFs, we performed scRNA-seq profile of CFs at 3 and 5 dpi. The novel data was integrated with previously reported scRNA-seq data[4] from healthy and 7 dpi CFs. A total of 27,995 CFs were grouped in 17 different cell clusters (Figure [C]). In healthy cardiac tissue, *Postn*^+^ CFs were mostly found in cluster 9. After MI, the number of these cells increased in almost every cluster (Figure [D]). In contrast, the first RCFs were observed after infarct in clusters 5, 8, and 11, and increased their number onwards (Figure [E]). These results suggest a progressive transition from early *Postn*^+^/*Cthcr1*^-^ activated CFs towards a *Postn*^*+*^*/Cthrc1*^*+*^ RCF profile (between 3 and 5 dpi) and are consistent with RCF’s healing role during scar formation.

To further characterize this transition, we investigated the ventricular distribution of *Postn*^+^/*Cthcr1*^-^ versus *Postn*^+^/*Cthcr1*^+^ using spatial transcriptomics. To achieve this goal in an unbiased manner, we first adapted a previously published criterion to identify anatomical domains based on cardiomyocyte transcriptomic profiles [5]. Our approach identifies four different domains on infarcted heart tissue samples: remote zone (RZ), border zone 1 (BZ1, closer to RZ), border zone 2 (BZ2, closer to the infarct zone/IZ), and IZ (Figure [F]). Such domains tightly correlate with the histomorphological traits observed after hematoxylin-eosin staining of the same tissue sections (Figure [F]). Our results show that both subpopulation signatures (*Postn*^+^/*Cthcr1*^-^ and *Postn*^+^/*Cthcr1*^+^) are enriched in the IZ at 3 dpi. At 5 dpi, however, this up-regulation is more specific for RCFs. Moreover, the RCF transcriptional signature is gradually enriched, following a characteristic gradient from the RZ to the IZ. These results strongly suggest a transcriptional shift from BZ activated CFs to IZ RCFs (Figure [F]).

To unravel the global transcriptional dynamic involved in activated CF to RCF transition, a ranking-based RNA velocity analysis per cell cluster was carried out using *velocito* [6]. Data from this analysis revealed that all cell clusters can be grouped into two dynamics (D1, D2), of which only D2 corresponds to RCFs (Figure [G]). D2 genes were associated with the “TGF-β signaling pathway”, and extracellular matrix molecules, such as *Fndc1, Ltbp3*, or *Aspn*. These results suggest that post-MI specification of RCFs critically depends on their interaction with the extracellular matrix/environment. The spatial location of D1 and D2 transcriptomic signatures indicates a clear enrichment of D2 genes in the IZ at both 3 and 5 dpi (Figure [H]). Altogether, our data suggest a crucial role of D2 genes in activating RCFs.

Among D2 genes, we have identified Asporin (*Aspn*) as a suitable requisite for RCF specification based on different criteria. First, *Asp* is an extracellular matrix (ECM) protein that acts as a natural inhibitor of the canonical “TGF-β signaling pathway”. Second, *Aspn* loss-of-function yields a *Cthcr1*^-/-^-like ventricular rupture phenotype at 5 dpi in mice [7]. Third, *Aspn* expression is clearly upregulated in post-MI CFs when compared with healthy ones in our analysis (Figure [I]). When *Aspn* expression was evaluated on infarcted heart tissues using RNAscope, we observed a gradual shift from *Postn*^*+*^*/Aspn*^*+*^*/Cthrc1*^*-*^ activated CFs, at the most external region of the BZ, to *Postn*^*+*^*/Aspn*^*-*^*/Cthrc1*^*+*^ RCFs, in the most internal area of the fibrosis (Figure [J]). Taken together, these results highlight the relevance of *Aspn* in the transition of activated CF to RCF.

To assess the translational potential of our findings, we analyzed the expression of *ASPN* in biopsies collected from infarcted pig hearts[4] (Figure [K]). We observed a temporal window of *ASPN* up-regulation in the infarcted ventricle (IZ), which correlates with the expression of both *POSTN* and *CTHRC1*. Finally, we studied the transcriptional profile of human biopsies samples collected from the RZ and IZ of patients with ischemic cardiomyopathy[4]. This approach identified a subset of D2 genes that are significantly upregulated within MI samples (Figure [L]), suggesting a role for D2 genes in the ventricular remodeling process in patients.

Together, our results narrowed down the activation point for RCFs, highlighting the relevance of the ventricular remodeling period in the healing fibrosis, and identifying an optimal therapeutic window for the management of the infarcted heart. This window is characterized by the expression of a gene set, including *ASPN*, which are directly involved in the induction of RCFs.

Although more investigation is required, these genes could be explored as molecular candidate targets for the control of fibrotic scar, avoiding clinical complications associated with increased scar size, including life-threatening arrhythmias and pump failure. Moreover, considering that a similar fibroblast subpopulation has been recently described in fibrotic processes in other organs [2, 3], the study of the mechanisms underlying the induction of this emergent population could offer new therapeutic strategies targeting a wide array of diseases characterized by fibrosis.

## Acknowledgments

The authors thank D.A. Brenner and T. Kisseleva (University of California San Diego) for the gift of Col1*α*1-GFP mice

