## Supplementary material for "Unravelling the molecular activation of the reparative cardiac fibroblasts after myocardial infarction": Material and Methods

### SUPPLEMENTAL METHODS

All data, analytic methods and study materials are available from the corresponding author upon request.

#### Animal models

All animal requisitions, housing, treatments and procedures were performed according to all state and institutional laws, guidelines and regulations. All studies were approved by the Ethics Committee for Animal Research at the University of Navarra and the Government of Navarra. *Collagen1α1-GFP* mice were previously described [1].

#### Induction of MI in mice

Myocardial infarction (MI) was induced in mice by ligation of the left anterior descending (LAD) coronary artery as previously described [2]. Briefly, 8 to 10 weeks old mice were anesthetized with vaporized isoflurane, intubated using a 20G intravenous catheter, mechanically ventilated, and placed on a heating pad to maintain body temperature. A left thoracotomy was performed at the fourth-fifth intercostal space, where muscles were dissected. The LAD coronary artery was permanently ligated using a 7/0 non-absorbable ethylene suture. After visual verification of anemia and akinesis of the apex and anterior-lateral wall to ensure coronary occlusion, the thorax was closed in layers. After extubation, mice were kept warm until fully recovered. Healthy or infarcted mice at 1 to 7 dpi were sacrificed and processed for analyses.

#### Human samples

The study protocol was approved by the Medical Ethics Committee and informed consent was obtained from all patients. Human heart samples were kindly provided by Professor

S. Janssens. Heart biopsies from the right (RV) and left ventricle (LV) of healthy donors (n=6), from patients with dilated cardiomyopathy (DCM) (n=5), and from infarct (IZ) and remote zone (RZ) of patients with a history of chronic MI (n=8) were collected for molecular analysis.

**Single cell optimization and preparation**

Mouse cardiac interstitial cells (CIC) were obtained from individual, 8-10 week-old mice as previously described [2]. Briefly, after euthanasia the thorax was opened and the heart was perfused with ice-cold phosphate buffered saline pH 7.6 (PBS) (Lonza), atria were excised and discarded, and both ventricles were digested completely. Excised tissues were placed in ice cold DMEM medium (Sigma) supplemented with 10% Fetal Bovine Serum (FBS) (Hyclone, GE). Ventricles were minced using a sterile scalpel. Pieces of tissue were incubated on an orbital shaker for 10 min at 37 °C in the presence of Liberase TH (125 μg/mL) (Roche) in HBSS++ solution (Hanks balanced salt solution, Gibco). After enzymatic incubation, partially digested tissue was mechanically dissociated by slowly pipetting to generate a single cell suspension. The supernatant was filtered through a cell strainer to discard cardiomyocytes (CM) (40 μm, nylon; Falcon). The digestion was repeated with the sedimented pieces, and the supernatants were pooled together. Erythrocytes were removed using RBC lysis buffer (eBioscience). The total time for enzymatic digestion was 30 min.

In order to include all cardiac fibroblasts (CF) in the following studies, an optimization step was performed as described previously [3]. Briefly, total CIC pellet was resuspended in 80 μL sorting buffer (2 mM EDTA, 0.5% BSA in PBS) and incubated with 20 μL of Feeder Removal MicroBeads (mEFSK4) (Miltenyi) for 15 min at 4 °C. A positive selection of CIC, enriched in CF, was performed twice using LS columns (Miltenyi) according to the manufacturer's instructions.

**Cell sorting**

The positive fraction from the LS column enrich in CF was centrifuged at 1,200 rpm and the pellet was resuspended in 100 µL of sorting buffer. Cells were incubated with the corresponding antibodies for 15 min at room temperature in the dark **(Table I in the Supplement)**. After incubation, the samples were washed twice with sorting buffer and spun at 1,200 rpm for 5 min, supernatant was discarded, and the final pellet was resuspended in 250 µL sorting buffer. Cell sorting was performed using FACSAria (BD Biosciences) and analyzed with FACSdiva software (BD Biosciences). Standard, strict forward scatter width *versus* area criteria were used to discriminate doublets and gate only singleton cells. Viable cells were identified by staining with 7-AAD (BD Bioscience). Viable cells gated on the FSC/SCC were sorted on the basis of the expression of GFP ($GFP^+/CD31^-/CD45^-$), and/or staining with anti-feeder cells antibody (mEFSK4 clone) (Miltenyi).

**Bulk MARS-seq**

Bulk RNA-seq was performed following a massively parallel RNA single-cell sequencing (MARS-seq) protocol adapted for bulk RNA-seq [4, 5] with minor modifications. Briefly, 3,000 to 10,000 cells were sorted in 100 µL of Lysis/Binding Buffer (Ambion), vortexed and stored at -80 °C until further processing. For cultured CF, samples were trypsinized, homogenized in 200 µL of Lysis/Binding Buffer and stored at -80 °C until further processing. Poly-A RNA was extracted with Dynabeads Oligo (dT) (Ambion) and reverse-transcribed with AffinityScript Multiple Temperature Reverse Transcriptase

(RT) (Agilent) using poly-dT oligo primers (IDT) carrying a 7 bp index. Up to 8 samples with similar overall RNA content were pooled together and subjected to linear amplification via IVT using the HiScribe T7 High Yield RNA Synthesis Kit from New England Biolabs (NEB). The resulting antisense RNA was fragmented into 250-350 bps fragments with RNA Fragmentation Reagents (Ambion) and dephosphorylated with 1U FastAP (Thermo Scientific) for 15 min at 37 °C. Partial Illumina adaptor sequences [4] were ligated with T4 RNA Ligase 1 (NEB) followed by a second reverse transcription. Full Illumina adaptor sequences were added during library amplification with KAPA HiFi DNA Polymerase (Kapa Biosystems). Libraries were quantified using a Qubit 3.0 Fluorometer and their size profiles examined in Agilent's 4200 TapeStation System. Sequencing was carried out in an Illumina NextSeq 500 using paired-end, dual-index sequencing (Read1: 68 cycles; Read2: 15 cycles; i7 index: 8 cycles) at a depth of 10 million reads per sample. Between 2 and 4 biological replicates per population were performed.

**Truseq RNA-sequencing**

Total RNA from frozen biopsies of human hearts was isolated using TRIzol reagent (Ambion). Following mechanical homogenization with an Ultra-turrax (T10 basis Ultra-Turrax, IKA), RNA was either extracted according to the manufacturer's instructions or stored at -80 °C until processed. RNA concentration was quantified using a Qubit 3.0 Fluorometer and its quality was examined in Agilent's 4200 TapeStation System. RNA was subjected to rRNA depletion using Truseq Stranded Total RNA library prep Gold kit (Illumina), followed by RT, second strand synthesis, 3' adenylation, Y shaped adaptor ligation and library enrichment as described above. Final libraries were quantified, and their profiles examined as described above. Sequencing was performed in an Illumina

NextSeq 500 using single-end, dual-index sequencing (Rd1: 75 cycles; i7: 8 cycles; i5: 8 cycles) at a minimum depth of 20 million reads per sample (n=21).

**Single-cell RNA-sequencing (scRNA-seq)**

The transcriptome of isolated $GFP^{+}$-CF from 8-12 weeks old, healthy or infarcted mice (3, 5 or 7 days post infarction (dpi)) were examined using Single Cell 3' Reagent Kits v2 (10X Genomics) according to the manufacturer's instructions. Two hearts were pooled in each scRNA-seq experiment. Briefly, 25,000 $GFP^{+}/CD31^{-}/CD45^{-}$ or $mEFSK4^{+}/CD31^{-}/CD45^{-}$ events were sorted in 1X PBS, 0.05% BSA and the number of cells was quantified in a Neubauer chamber. Approximately 16,000 cells were loaded at a concentration of 1,000 cells/µL on a Chromium Controller instrument (10X Genomics) to generate single-cell gel bead-in-emulsions (GEMs). In this step, each cell was encapsulated with primers containing a fixed Illumina Read 1 sequence, followed by a cell-identifying 16 nt 10X barcode, a 10 nt Unique Molecular Identifier (UMI) and a poly-dT sequence. Upon cell lysis, reverse transcription yielded full-length, barcoded cDNA. This cDNA was then released from the GEMs, PCR-amplified and purified with magnetic beads (SPRIselect, Beckman Coulter). Enzymatic Fragmentation and Size Selection was used to optimize cDNA size prior to library construction. Fragmented cDNA was then end-repaired, A-tailed and ligated to Illumina adaptors. A final PCR-amplification with barcoded primers allowed sample indexing. Library quality control and quantification was performed using Qubit 3.0 Fluorometer (Life Technologies) and Agilent's 4200 TapeStation System (Agilent), respectively. Sequencing was performed in a NextSeq500 (Illumina) (Read1: 26 cycles; Read2: 57 cycles; i7 index: 8 cycles) at an average depth of 50,000 reads/cell.

**Tissue processing and Visium data generation**

The hearts from *Collagen1α1-GFP* 8-12 weeks old, healthy or infarcted mice (3 and 5 dpi) were excised and washed in PBS, embedded in OCT compound, frozen in dry ice and sectioned in a cryostat at a thickness of 10 µm at -20 °C following the manufacturer's recommendations.

In order to check the RNA quality of the tissues, ten tissue sections of each sample were collected in an RNase-free Eppendorf tube, and RNA was extracted with the Qiagen RNeasy extraction kit (Qiagen) according to the manufacturer's instructions. RNA quality was assessed with a 4200 TapeStation (Agilent Technologies), and all the samples had a RIN between 7.2 and 7.7.

For optimization of the tissue permeabilization, seven sections from the 3 dpi sample were collected onto a -20 °C pre-equilibrated Visium tissue optimization slide (10X Genomics) as per the manufacturer's recommendations. Briefly, tissue sections were fixed in chilled methanol, then stained with Mayer's Hematoxylin solution (Millipore Sigma), followed by Bluing Buffer (Dako), and Eosin Y solution (Millipore Sigma). Hematoxylins & Eosin (H&E) stained tissues were imaged on an Aperio CS2 (Leica Biosystems) digital slide scanner at 40X magnification. Tissues on optimization slides were permeabilized in a time course experiment from 3 to 30 minutes, and reverse transcription was performed using fluorescently labeled nucleotides, resulting in fluorescent cDNA bound to the capture areas within the slide. Tissues were enzymatically removed, and fluorescence imaging was performed using a Zeiss LSM800 (Carl Zeiss Microscopy) digital slide scanner with a Red filter set. Whole slide scan was carried out using a 20X magnification objective and with 150 milliseconds exposure per image frame. Sequentially imaged frames were automatically stitched. A permeabilization time of 12 minutes resulted in the maximum fluorescence signal with the lowest signal diffusion and was chosen as the optimal time to perform the subsequent Visium expression experiments.

In the final experiment, the mRNA capture was performed following instructions of the Visium Spatial Gene Expression Reagent Kits User Guide (10x Genomics). Briefly, one section of each sample was collected onto a pre-equilibrated Visium expression slide. After H&E staining, stained tissues were imaged on the Aperio CS2 (Leica Biosystems) digital slide scanner at 40X magnification. The sections were permeabilized for 12 minutes to release poly-adenylated mRNA from overlying cells onto the capture areas of the slide. The bound mRNAs were then reverse transcribed, resulting in spatially barcoded, full-length cDNA. Second strand synthesis was performed, followed by denaturation and transfer of the cDNA from the slide to a PCR tube. qPCR (KAPA SYBR FAST qPCR Master Mix, KAPA Biosystems) was used to determine the number of cDNA amplification cycles required based on the 25% RFU peak. After 16 cDNA amplification cycles, fragment analysis of the cDNA was performed on a 4200 TapeStation (Agilent Technologies). Library construction, performed on 25% of the cDNA, consisted of enzymatic fragmentation of the cDNA, end-repair, A-tailing and adaptor ligation, followed by a sample index PCR. Final libraries containing P5 and P7 primers were quantified with Qubit 3.0 Fluorometer (Life Technologies). Libraries were loaded at 650 pM and sequenced on a Nextseq 2000 System (Illumina) using a Nextseq2000 P3 Reagent Kit (100 cycles, Illumina), at an average sequencing depth of approximately 100,000 reads per spot (25-35% of the 5,000 total spots on the slide were used). Sequencing was performed using the following read protocol: Read1, 28 cycles; i7 index, 10 cycles; i5 index, 10 cycles; Read2, 91 cycles.

**Immunofluorescence**

Mice adult hearts were excised and washed in PBS, fixed in 4% fresh paraformaldehyde (Sigma). Some specimens were then cryoprotected in sucrose (Sigma), embedded in OCT compound (VWR), frozen in dry ice and stored until sectioning. For Reparative Cardiac Fibroblast (RCF) specific marker staining additional specimens were dehydrated, paraffin embedded, sectioned and rehydrated before the staining. Five-to-ten µm thick sections were washed in Tris-PBS (TPBS), non-specific IgG binding sites were blocked with 8% goat serum (Dako), 1% BSA, and 0.1% Triton X-100 (Sigma) followed by incubation overnight with the corresponding primary antibodies at 4 °C **(Table I in the Supplement)**. Anti-GFP staining required an antigen retrieval step using 100 mM citrate buffer pH6.3 (Sigma) before blocking. The sections were then washed and incubated with the appropriate fluorescence-conjugated secondary antibodies for 1 hour before mounting **(Table I in the Supplement)**. Negative controls were included by omitting the primary antibody. Cell nuclei were counterstained with 4',6-diamidino-2-phenylindole (DAPI, Sigma). All images were captured in a Zeiss LSM 510 800 (Zeiss) or Leica SP5 confocal laser microcopy (Leica).

**RNA *in situ* hybridization (ISH) assay by RNAscope®**

Mice adult hearts were fixed in formalin overnight at 4 °C and washed three times with PBS for 5 min. Then, samples were dehydrated, paraffin embedded, and sectioned into 10 µm slides. Sections were analyzed with an RNAscope® assay, using the RNAscope® Multiplex Fluorescent Reagent Kit v2 (Advance Cell Diagnostics) following manufacturer's instructions. Briefly, sections were dehydrated, deparaffinized, and incubated with hydrogen peroxide solution. Subsequently, sections were treated with RNAscope® Target Retrieval reagent solution, and then protease plus was applied in a humid chamber. Then, sections were incubated with the following targeted probes: *Postn*, *Aspn*, *Cthrc1*, and RNAscope™ 3-plex Negative Control probe (all from Advance Cell Diagnostics). The hybridization procedure was performed for 2 h at 40 °C, followed by an overnight incubation with 5X SSC buffer (20X SSC Buffer, Invitrogen) at room temperature. The next day, sections were incubated with the detection reagents for manual amplification and washed twice with the RNAscope® Wash Buffer. After amplification, sections were incubated subsequently with the appropriate HRP channel, the corresponding Opal (Akoya Bioscience) and the HRP blocker, with RNAscope® Wash Buffer between incubations. Finally, the sections were incubated with DAPI and mounted with ProLong™ Gold antifade reagent (Invitrogen).

**Bulk RNA-seq analysis**

Samples were demultiplexed using Illumina *bcl2fastq* software (1.2.4) and aligned with the mouse (*mm10*) or human (*GRCh38*) genome with STAR (2.6.1)[6] setting the parameters to default values. Quantification and generation of gene expression matrices were performed with the function *featureCounts,* implemented in the R [7] package *Rsubread* [8]. The *ensembl* transcriptomes (*GRCm38.91* and *GRCh38.92*) were used as reference for gene annotation. An initial filtering of not expressed genes was performed. Additional filtering, data transformation, normalization, and testing for differential expression was performed with DESeq2 [9].

For human samples, GSVA [10] was used to compute scores for the expression of different gene signatures. We applied GSVA to the DESeq2 normalized expression matrix. We calculated scores for the following signatures: 1) Velocity signature 2) gene cluster 4 3) gene cluster 9 4) the ratio between gene cluster 4 and 9.

**Single cell RNA-seq analysis**

Sequenced libraries were demultiplexed, aligned to the mouse transcriptome (*mm10*) and quantified using *Cell Ranger* (3.0.1) from 10X Genomics. The output of the pre-processing pipeline consisted of gene expression matrices per cell. Further computational analysis was performed using *Seurat* (3.1.0). Cells were subjected to quality control filters based on the number of detected genes, number of UMIs and proportion of UMIs mapped to mitochondrial genes per cell. Regarding the number of genes and UMI, we established the first and fourth quartile as lower and upper threshold, respectively. Cells with > 5% mitochondrial genes were also filtered out. Using these parameters, 6,277 (healthy myocardium), 7,888 (3 dpi), 4,839 (5 dpi), 8,991 (7 dpi) cells were retained.

Each single cell dataset was subjected to log normalization and scaling. Additional normalization and variance stabilization, together with removal of unwanted source of variation (% of mitochondrial genes and UMI numbers), was performed using the SCTransform method [11].

Next, we integrated the 4 time points using a set of anchors that was found based on canonical correlation analysis (CCA) [12]. A total of 2,000 variable features were used as input for the analysis and 30 canonical vectors were calculated.

The integrated single cell dataset was rescaled and subjected to principal component analysis (PCA). Unsupervised clustering was performed, using the 20 first PCs and a resolution parameter of 0.8. Non-linear dimensional reduction was computed using uniform manifold approximation and projection (UMAP) [13].

Gene signature characterizing RCFs was obtained from Ruiz-Villalba *et al* [3]. We selected the set of markers from cluster B with logFC > 0.1 and adjusted p value < 0.01. This signature was used as input for AUCell [14], in order to explore its activity in the individual cells. Expression ranking was created with the normalized counts (after SCTransform).

All differential expression tests were done using the function *FindAllMarkers* with the MAST test [15].

**RNA Velocity**

For the velocity analysis, we created a subset of the Seurat dataset selecting exclusively the cells expressing *Postn*. Every cell displaying *Postn* raw counts > 0 was considered *Postn*$^{+}$ and included in the subset. We reprocessed this subset following the steps described above (log normalization, scaling, SCTransform and integration) and computed a new UMAP. After reprocessing, we used the new SCT corrected matrix as input for the velocity.

The spliced/unspliced matrices were generated with *velocyto* [16] (0.17.17) for each condition (healthy, 3, 5 and 7 dpi) using the BAM files from *CellRanger* (3.0.1) as input. Further processing was performed using *scVelo*[17] (version 0.2.3) implemented in *Python* (3.8.8). Data was preprocessed by filtering the top 2000 highly variable genes, PCA and neighborhood graph construction, using default options. The dynamical model was used to estimate RNA velocities. The stream plots were projected on the UMAP reductions retrieved with *Seurat*. Differential expression analysis was applied to the gene velocities, using the function *rank_velocity_genes,* to rank genes based on differential dynamics across cell clusters. Genes resulting from this analysis were finally subjected to hierarchical clustering using their ranking positions as input values.

**Spatial gene expression analysis**

Visium libraries were demultiplexed and mapped with *Space Ranger* (1.3.1) from 10X Genomics. Libraries were mapped to an Ensembl 105 based reference (GRCh39 reference, v3.0.0). The reference was modified to include the GFP gene, following 10X instructions. Manual alignment was performed with *Loupe Browser* (6.1.0) to select spots under the tissue. Spatial transcriptomic data was processed using *Seurat* [12] (4.1.1). Quality control was performed in order to remove spots with low quality defined as UMI < 500, gene < 250 and mitochondrial (%) > 50. After, we normalized data using the regularized negative binomial regression implemented in the *sctranform* [11] R-package and a visual exploration was performed through PCA, unsupervised shared-nearest-neighbor-based clustering and UMAP. Enrichment analysis for predefined signatures (cluster B/RCFs, gene cluster 4 and 9 signatures) was performed using *AUCell* [14] (1.16.0) for each spot in the samples. After this step, we adapted a published criterion [18] to identify anatomical domains based on cardiomyocyte transcriptomic profiles, dividing the heart tissue samples spots on remote zone (RZ), border zones 1 (BZ1, closer to RZ), border zone 2 (BZ2, closer to the infarct zone/IZ), and infarct zone (IZ). Enrichment score was then plotted by segment to understand the score gradient in the RCFs, which included cluster 5, 8, 9 and 11, and the score in dynamics 1 and 2 and the ratio between them. Linear regression was performed per spot with *stats* [7] (3.6.2) to study the dependence of the signature's gradient with the fibroblast population in the different areas of the tissue.

**Data resources**

All single-cell RNA-seq and bulk RNA-seq from previous the previous study, data are available in a SuperSeries at NBCI's Sequence Read Archive database under accession number GSE132146. All single-cell RNA-seq data from experiments performed for this study are available at NBCI's Sequence Read Archive database under accession number GSE261428. New bulk RNA-seq added is available under accession number GSE267256. All spatial transcriptomics data are available in a SuperSeries at NBCI's Sequence Read Archive database under accession number GSE265828.

**Supplementary Table I: List of antibodies used.**

| **Name** | **Clone** | **Supplier** | **Dilution** | **Use/Technique** |
|---|---|---|---|---|
| CD31 (PECAM)-APC | MEC 13.3 | BD Pharmingen, 551262 | 1:200 | FACS |
| CD45-PE | 30-F11 | eBioscience 12-0451-81 | 1:200 | FACS |
| Anti-feeder cells-APC | mEFSK4 | Miltenyi, 130-102-302 | 1:50 | FACS |
| CTHRC1 | Vli55 | Maine Medical Ctr. Res. Institute | 100 ng/ml | IF |
| GFP | pAb | Abcam, ab13970 | 1:100 | IF |
| Anti-Chicken 488 | 2ry Ab | Jackson Inmunoresearch, 703-545-155 | 1:200 | IF |
| Anti-Rabbit 647 | 2ry Ab | Invitrogen, A31573 | 1:200 | IF |
